# Endometrial transcriptomic and ciliation analysis after scratch shows no signature linked to live birth following IVF

**DOI:** 10.64898/2026.07.20.739558

**Authors:** Ella Proudley, Jennifer Pearson-Farr, Ian Reddin, Clare Pye, Susan Laird, Rohan M Lewis, Jane K Cleal, Mostafa Metwally, Ying Cheong

## Abstract

**Objective:** To determine whether endometrial scratch induces differences in transcriptomic profiles or epithelial cell ciliation in the endometrium at the window of implantation that associate with live birth following IVF.

**Design:** Secondary analysis of 50 matched endometrial biopsies collected within a randomised controlled trial evaluating the clinical effectiveness of endometrial scratch before first-time IVF.

**Setting:** Endometrial biopsy samples were obtained from women attending the Jessop Wing of Sheffield Teaching Hospitals.

**Population or Sample:** Women undergoing first-time IVF who received an endometrial scratch in the preceding cycle at the window of implantation (6–10 days after LH surge).

**Methods:** Endometrial biopsies were molecularly dated using transcriptomic menstrual-cycle staging algorithms. Bulk RNA sequencing was analysed using DESeq2 with FDR correction, and principal component analysis (PCA) assessed clustering patterns. Epithelial cell ciliation was quantified using immunohistochemistry and an automated Python-based image analysis pipeline.

**Main Outcome Measures:** Differential endometrial gene expression between women with and without live birth after IVF; percentage coverage of ciliated epithelial cells in luminal and glandular regions.

**Results:** Transcriptomic dating confirmed no differences in menstrual-cycle timing between live-birth and no-live-birth groups. No significant differential gene expression was detected (log2FC >2, FDR <0.05), and PCA showed no clustering by pregnancy outcome. Ciliation coverage did not differ between outcome groups or between glandular and luminal surfaces.

**Conclusions:** When implantation timing is precisely defined, endometrial scratch does not produce detectable transcriptomic changes or alterations in epithelial ciliation that distinguish women who achieve live birth after IVF.

**Funding:** Wellbeing of Women RG2147; Wessex Medical Research; Rosetrees Trust (PGS23/100171).

## Introduction

Implantation remains the principal rate-limiting step in in vitro fertilisation (IVF). Despite advances in embryo culture, selection, and laboratory practice, live birth rates per embryo transfer plateau at approximately 30–35%, and a substantial proportion of IVF failure remains unexplained. While IVF can overcome tubal and male-factor infertility, endometrial competence remains intrinsic to reproductive success. This has driven sustained interest in interventions targeting endometrial receptivity, particularly in women experiencing implantation failure.

Endometrial scratch (ES), an intentional mechanical injury to the endometrium, typically performed with a Pipelle catheter, was introduced after early observational data suggesting improved implantation after local injury (Barash et al., 2003). Proposed mechanisms include induction of a wound-healing inflammatory cascade, altered cytokine signalling, enhanced decidualisation, modulation of receptivity genes, and improved embryo–endometrial crosstalk (Gnainsky et al., 2010, Li and Hao, 2009). These biologically plausible hypotheses led to the rapid and widespread adoption of ES as an IVF “add-on”, often before robust mechanistic or clinical validation.

High-quality randomised trials have since challenged its effectiveness. In the large multicentre UK Endometrial Scratch Trial, ES performed in the mid-luteal phase of the menstrual cycle prior to first-time IVF did not improve live birth rate (Metwally et al., 2021). Similarly, systematic reviews focused on first-cycle IVF have shown no convincing benefit of ES (Metwally et al., 2022, Vitagliano et al., 2018). Despite this, ES continues to be offered in some settings, a phenomenon attributed to cognitive dissonance and persistence of disproven interventions within reproductive medicine (Paulson, 2022).

Conversely, some analyses have suggested potential benefit of ES in selected subgroups. A systematic review in women with one or more previous failed embryo transfers reported higher live birth rates following ES (Vitagliano et al., 2018). More recently, an individual participant data meta-analysis reported a modest overall increase in live birth following ES (van Hoogenhuijze et al., 2023), although there was no clear interaction with age, infertility cause, or number of previous failures. A broader meta-analysis has similarly suggested improved pregnancy outcomes following ES (Maged et al., 2023). However, these findings are characterised by substantial heterogeneity across patient populations, injury timing, techniques, and trial quality.

Crucially, the biological premise underpinning ES remains incompletely established. The endometrium undergoes tightly regulated transcriptomic remodelling across the menstrual cycle, with a distinct molecular signature during the window of implantation. Advances in bulk and single-cell RNA sequencing have refined our understanding of receptivity-associated gene expression patterns and cellular heterogeneity. However, whether mechanical endometrial injury induces sustained transcriptomic changes in a subsequent cycle remains uncertain. In a longitudinal study examining paired luteinising hormone (LH) +7 days endometrial biopsies before and after mid-luteal ES in women with recurrent implantation failure, no significant differences were observed in whole-genome transcriptomic profiles or established receptivity gene sets (Tian et al., 2023). These data raise the question of whether ES exerts a durable molecular effect at the time of implantation.

Beyond gene expression, the structural composition of the epithelium may influence implantation. The luminal epithelium comprises secretory and ciliated cell populations whose distribution varies across the cycle. Although ciliated cells are traditionally associated with tubal transport, they are present within the endometrium and may influence local fluid dynamics, embryo apposition, and epithelial receptivity. The relationship between endometrial epithelial cell ciliation and IVF outcome has not been systematically explored.

An additional methodological challenge in endometrial research is accurately dating the menstrual cycle. Reliance on menstrual history or urinary LH surge testing may introduce temporal misclassification within the narrow implantation window. Transcriptomic dating algorithms now allow molecular staging of biopsies, improving precision in comparative analyses. This study aims to determine whether endometrial scratch induces detectable differences in global transcriptomic profiles or menstrual cycle timing at the window of implantation in women undergoing first-time IVF and to assess whether epithelial ciliation coverage differs between women who achieve live birth following IVF and those who do not.

## Methods

### Data Collection and Ethical Approval

The endometrial biopsy samples used in this study were collected from women attending the Jessop Wing of Sheffield Teaching Hospitals. Ethical approval was granted by South Central Berkshire Research Ethics Committee (16/SC/0151), and the trial was registered with ISRCTN (ISRCTN23800982). The authors assume responsibility for the accuracy and completeness of the data analyses, adherence to the protocol, and interpretation of results.

### Sample Collection

The endometrial biopsy samples used in this study were collected from women attending the Jessop Wing of Sheffield Teaching Hospitals, as part of the clinical trial investigating the clinical effectiveness and safety of the endometrial scratch procedure prior to first-time IVF (Metwally et al., 2021). Ethical Committee approval for the whole clinical trial and the sub-study, which involved the collection and storage of tissue, was granted by South Central Berkshire Research Ethics Committee (16/SC/0151).

Participants were women aged 18–37□years (inclusive) who undergoing their first cycle of IVF, had a regular ovulatory menstrual cycle defined by clinical judgement or with ovulatory levels of midluteal serum progesterone, normal uterine cavity assessed by transvaginal sonography at screening, with no endometrial abnormalities that would require treatment to facilitate pregnancy (such as suspected intrauterine adhesions, uterine septum, submucosal fibroids or intramural fibroids exceeding 4□cm in diameter), good ovarian reserve assessed clinically, biochemically (FSH < 10 UI/L) and normal follicular phase oestradiol levels and/or normal anti-Müllerian hormone levels or sonographically (antral follicle count) and no history of previous radiotherapy or chemotherapy; had no relevant vaginal/uterine infections. Participants were excluded if they had received previous trauma to the endometrium (resection of uterine septum, intrauterine adhesions, or recent resection of significant submucous fibroids), had a BMI of 35 Kg/m2 or greater, were participating in another interventional fertility study, or had grade 4 endometriosis.

The women allocated to the intervention arm received the ES procedure in the mid-luteal phase (defined as 5–7□days before the expected next period, or 7–9□days after a positive ovulation test) of the menstrual cycle preceding IVF/ICSI by a suitably qualified doctor or nurse. ES was performed by inserting a speculum into the vagina, and the cervix was exposed and cleaned. A pipelle sampler or similar device was then inserted into the uterine cavity, and negative pressure was applied by withdrawing the plunger. The sampler was then rotated and withdrawn 3 to 4 times to allow tissue to appear in the transparent tube. The sampler and speculum were then removed.

The tissue from the sampler was divided into two parts and placed either in RNA later^TM^ (Ambicon) or 10% Neutral buffered formalin. The sample placed in RNA later was immediately frozen in liquid nitrogen and then stored at -80 C until use. The other sample was stored in 10% neutral buffered formalin at room temperature for 24 hours, after which it was passed through a series of alcohols (50%, 75%, 90% and 95%) and xylene before being embedded in paraffin wax blocks.

### Transcriptomic Endometrial Biopsy Dating

The R Package endest (Teh et al., 2023) was used to generate predicted values for each stage of the menstrual cycle (as a percentage of the menstrual cycle starting on day 1) based on RNA sequencing read counts. As the window of implantation is believed to occur during 70-80% of the cycle, samples dated outside this range were removed from the analysis, thereby eliminating the possibility that the observed differential expression is due to differences in menstrual cycle stage. A Welch’s test was performed to confirm whether there was a significant difference between the endometrial dating of samples taken from those who got pregnant after the scratch procedure and those who did not become pregnant.

### RNA extraction, library preparation and RNA sequencing

Total RNA was isolated from biopsy samples using the New England Biolabs Monarch Total RNA Miniprep Kit (NEB #T2010) according to the manufacturer’s instructions.

RNA sequencing (RNA-seq) libraries were prepared by Novogene UK Ltd (Directional mRNA enrichment libraries) and subjected to NovaSeq paired-end sequencing (2 × 150 base pairs) at an average coverage of 40 million total reads per library. Paired FASTQ files were aligned to the human genome using STAR 2.7.11b with the human genome 38 (GRCh38.p14) reference.

### Differential Gene Analysis

Paired FASTQ files were aligned to the human genome using STAR 2.7.11b with the human genome 38 (GRCh38.p14) reference. Gene count data were normalised, and differential gene expression was analysed in RStudio (R version 4.4.1) using DESeq2 (version 1.4.4.0). The empirical Bayes approach to false discovery rate (FDR) was used to adjust for multiple comparisons at the 0.05 significance level. Significance was assessed using a Wald test and considered statistically significant at p ≤ 0.05. A log fold-change threshold of 2 was applied to identify substantial changes in gene expression.

### Principal Component Analysis

To identify patterns in the data, principal component analysis (PCA) was performed using the prcomp function in RStudio (R version 4.4.1). The PCA was conducted using the default prcomp setting, which employs singular value decomposition (SVD). The resulting principal components were visualised in a biplot to assess clustering patterns and the variables’ contributions to each component.

### Immunohistochemistry

Endometrial biopsy sections embedded in paraffin were cut into 4 µm sections. Sections were washed twice in Dulbecco’s phosphate-buffered saline (PBS) for 5 minutes, followed by two 20-minute washes in PBS containing 0.05% BSA and 0.1% Triton X-100. They are then incubated overnight at 4°C in PBS with 0.05% BSA, 0.1% *Triton X-100*, and primary antibodies. Following three 10-minute washes in PBS with 0.05% BSA, 0.1% Triton X-100, samples are incubated at room temperature for 2 hours with secondary antibodies in 0.05% BSA, 0.1% Triton X-100, PBS, protected from light. After two 5-minute washes in PBS, then the sections are washed in 0.05% BSA, 0.1% Triton X-100, PBS containing a 1:500 dilution of DAPI for 20 minutes. The sample is mounted into a chamber slide following two 10-minute washes in PBS,

To investigate the coverage of multi-ciliated epithelial cells on both the glandular and luminal epithelia, the tissues were stained with Invitrogen™ Cytokeratin 19 Monoclonal Antibody (A53-B/A2.26 (Ks19.1)) (1:100), Invitrogen™ RSPH4A Polyclonal Antibody (1:250), and Invitrogen™ DAPI (4′,6-diamidino-2-phenylindole) (1:500). Each section was then imaged using a Leica Stellaris microscope. Five luminal regions and glandular regions of the biopsies were selected using the LAS X spiral navigator.

### Automated quantification of ciliated cells

Ciliated cell counts were obtained using a custom Python pipeline built with *scikit-image*. DAPI, Cytokeratin 19, and RSPH4A TIFF images were imported and pre-processed using Gaussian smoothing and intensity rescaling. Nuclei were segmented from the DAPI channel using Otsu thresholding and size-based filtering. Cytokeratin-19 and RSPH4A signals were thresholded using Otsu’s method with an additional upward bias to ensure conservative marker detection. For each labelled nucleus, spatial overlap with the Cytokeratin-19 and RSPH4A masks was assessed, enabling automated quantification of total epithelial cells and the subset that is also positive for RSPH4A. Automatic counting was validated by manually segmenting and counting 16 TIFF images in ImageJ. The Wilcoxon test showed no significant difference in counts between automated and manual counts of ciliated epithelial cells (p = 0.83, n = 32).

### Statistical analysis

A Welch’s test was performed to confirm whether there was a significant difference between the endometrial dating of samples taken from those who got pregnant after the scratch procedure and those who did not become pregnant. Non-parametric Mann Whitney test was performed to confirm whether there was a significant difference between the live birth and no live birth cohorts when investigating epithelial cell ciliation levels. For all statistical analyses, a p-value greater than 0.05 was deemed non-significant unless otherwise stated.

## Results

### Luteinising hormone surge urine tests, and endometrial transcriptomic dating methods effectively predict the window of implantation

The endometrial biopsies were dated (day of menstrual cycle defined) using two computational methods: one based on transcriptomic results and the other on a urine test to detect the LH spike at ovulation (day 14). Endometrial biopsy dating using the transcriptomic endometrial biopsy dating method(Teh et al., 2023) and the LH uterine test method both showed no significant difference between those who had a live birth after IVF and those who did not ***(Figure 1)***. This indicates that they were all correctly timed to each other within a few days and showing no evidence of cycle changes between the two groups.

**Figure 1:**
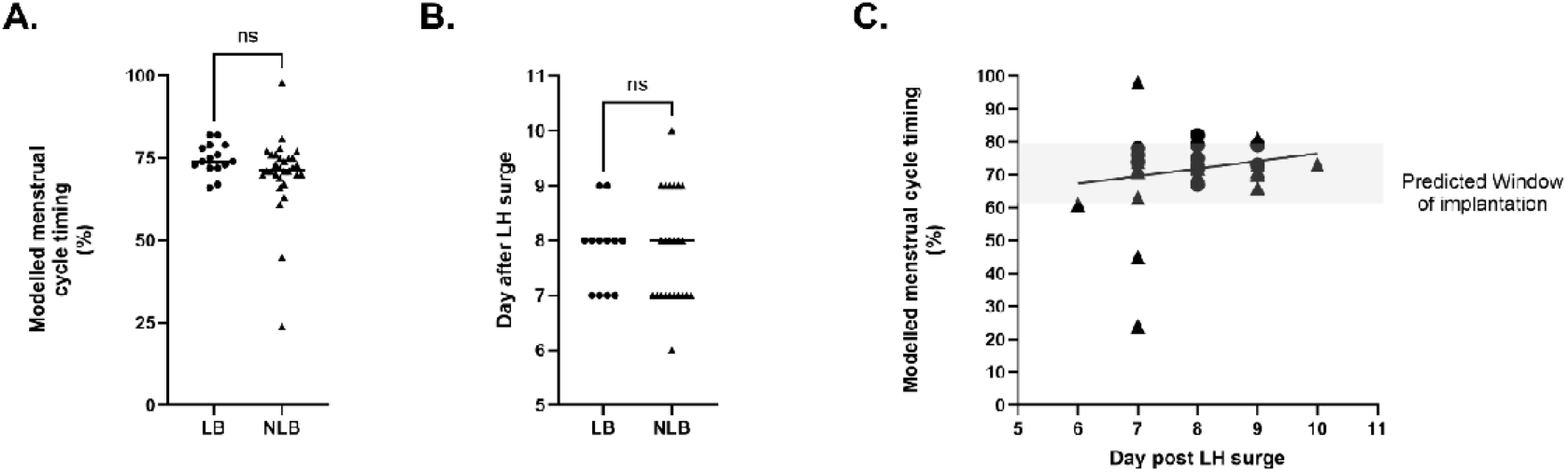
Endometrial biopsies were dated to confirm the stage/day of the menstrual cycle. LB = Live birth, NLB No live birth. A. Read count menstrual cycle dating shows no significant difference between the times samples were collected between participants who had a live birth after the scratch and those who did not. B. Luteinising hormone (LH) dating shows no significant difference between the days samples were collected in participants who had a live birth after the scratch and those who did not. Samples were collected 6 to 10 days after a recorded LH surge (n = 45, p = 0.08). C. Correlation of LH surge, menstrual dating, and modelled menstrual dating using RNA sequencing. LH surge testing reliably predicts the window of implantation. The shaded section shows the window of implantation recognised in the modelled menstrual cycle timing. **Ns = not significant**

### Differential gene expression shows no significant differences between those who had a live birth and those who did not after IVF

After confirming there were no endometrial dating differences between the groups, differential gene expression analysis was performed using the following restriction criteria: log2 fold change > 2 and FDR < 0.05 (p > 0.05, n = 45). When these restrictions were applied to the RNA sequencing data, the analysis revealed no significant difference in gene expression between those with a live birth and those without after an endometrial scratch. Furthermore, principal component analysis revealed no apparent clusters among the samples ***(Figure 2)***.

**Figure 2:**
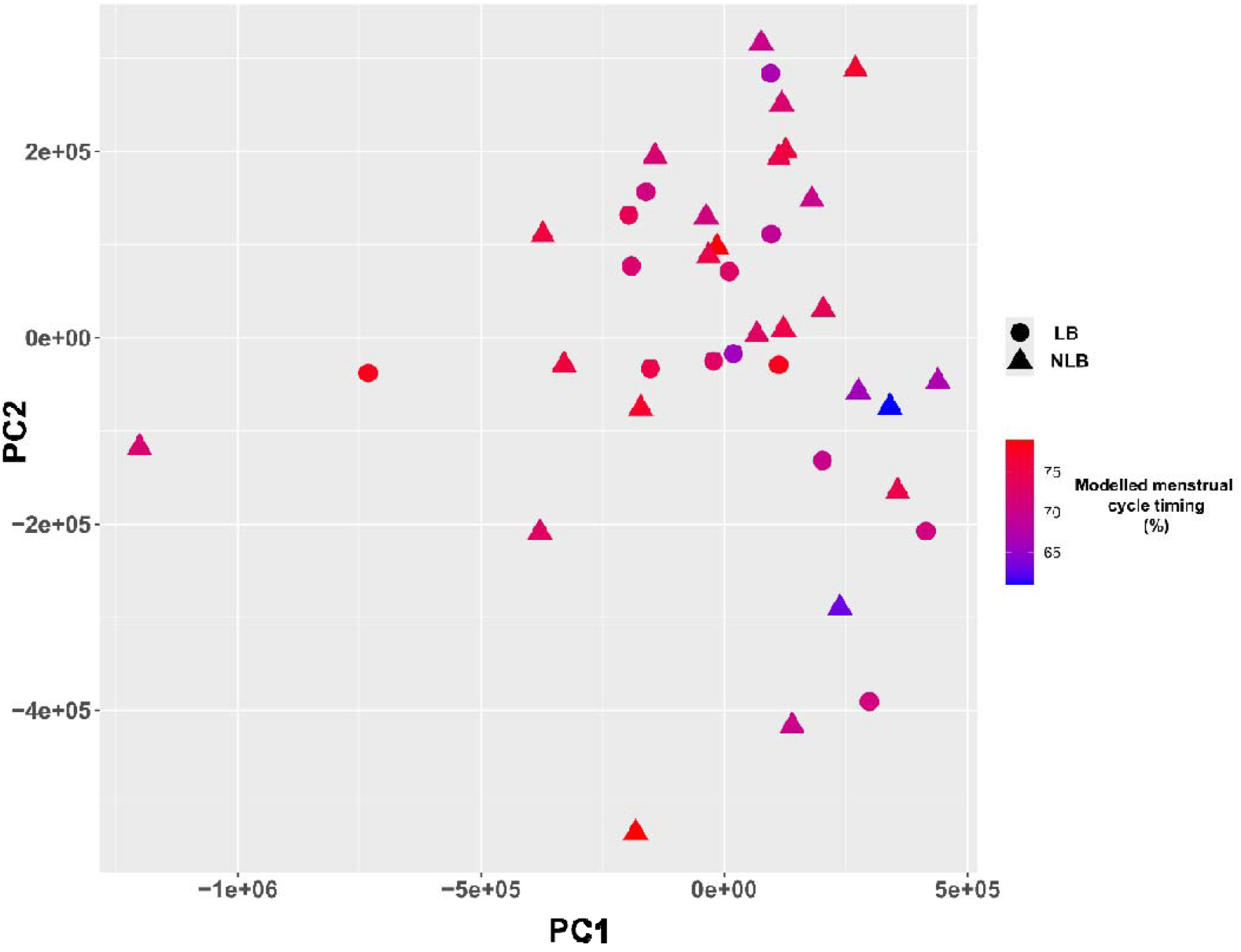
Principal component analysis (PC) shows no common endometrial transcriptional differences between those who had a live birth and those who did not after intervention through IVF and endometrial scratch. Log2 fold change > 2 and FDR < 0.05 (n = 45, p > 0.05). LB = live birth after IVF and scratch procedure. NLB = No live birth after IVF and scratch procedure. Menstrual cycle timing was predicted using bulk RNA sequencing analysis.

### There is no significant difference between the uterine luminal epithelial ciliation and the glandular epithelial ciliation

Immunohistochemistry and confocal imaging were used to calculate the percentage coverage of ciliated epithelial cells on fixed endometrial tissue sections. Five regions of both the glandular and luminal epithelium were imaged, and the numbers of ciliated and total epithelial cells were counted using scikit-image for image processing in Python. At the window of implantation, the average coverage of ciliated cells on the glandular epithelium as 1%, and the luminal epithelium was 4%. There was no significant difference in cilia coverage between the luminal and glandular epithelial (p = 0.206, n = 38; **Figure 3**).

**Figure 3:**
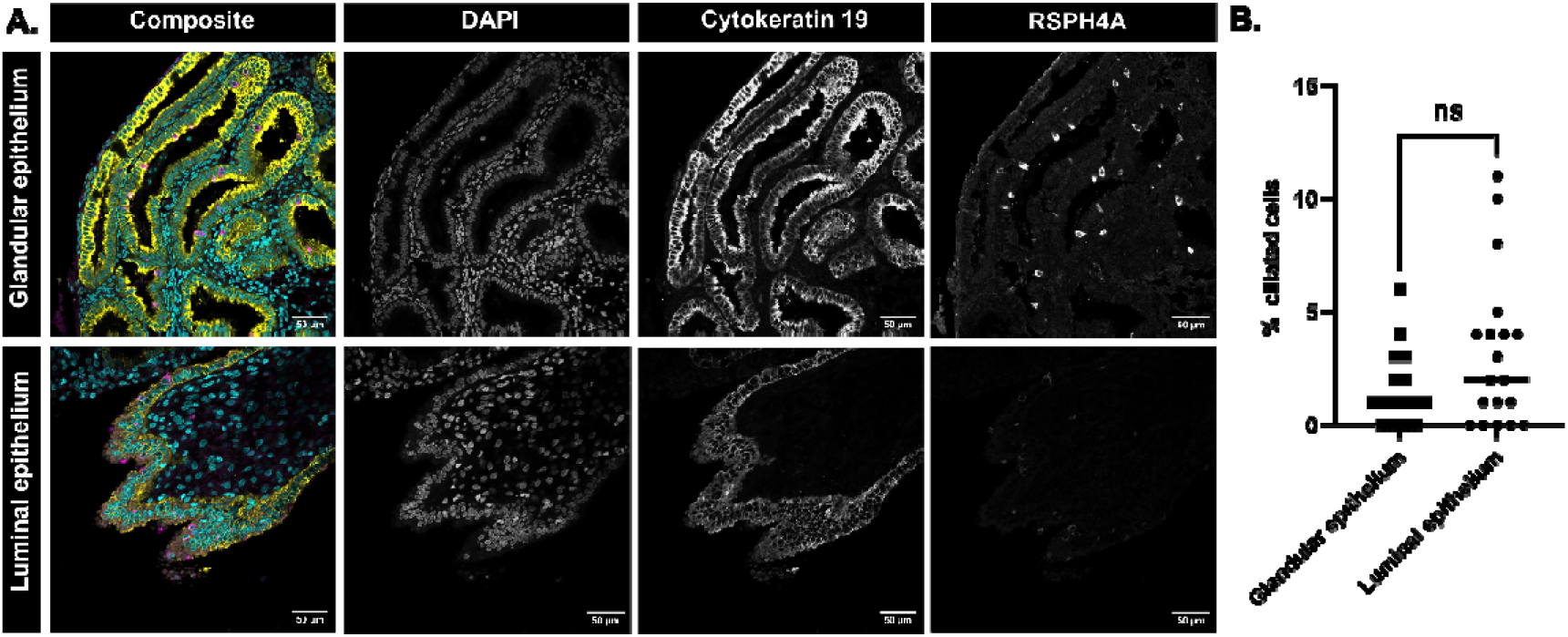
There is no significant difference between ciliation levels in the glandular and luminal endometrial epithelial surface at the window of implantation. **A.** Confocal micrographs of endometrial glandular and luminal epithelium at the window of implantation. DAPI staining nuclei (Cyan), Cytokeratin 19 staining epithelium (Yellow) and RSPH4A staining ciliated epithelial cells (Magenta). Scale bars = 50 µm **B**. There is no significant difference (ns) between the percentage of ciliated cells on the glandular epithelium and the luminal epithelium (n =38, p = 0.09). All samples are taken from the window of implantation. Percentage ciliation coverage on the glandular epithelium is 1%, and on the luminal epithelium is 4%. This is lower than the airway tissue, where motile cilia assist in mucus clearance.

### Ciliation levels show no significant differences between those who had a live birth and those who did not after

Further analysis of immunohistochemistry of the endometrial biopsies showed no significant difference between those who had live-birth and those who did not after endometrial scratch and IVF (p = 0.57, n = 44); ***Figure 4A, Figure 4B***). To investigate whether the distribution of ciliation differed between those who had live birth and those who did not, after endometrial scratch and IVF luminal ciliation was subtracted from glandular ciliation. This showed no significant difference in the distribution of cilia on the luminal and glandular surfaces between the two outcome cohorts (p = 0.82, n = 42;***Figure 4C***).

**Figure 4:**
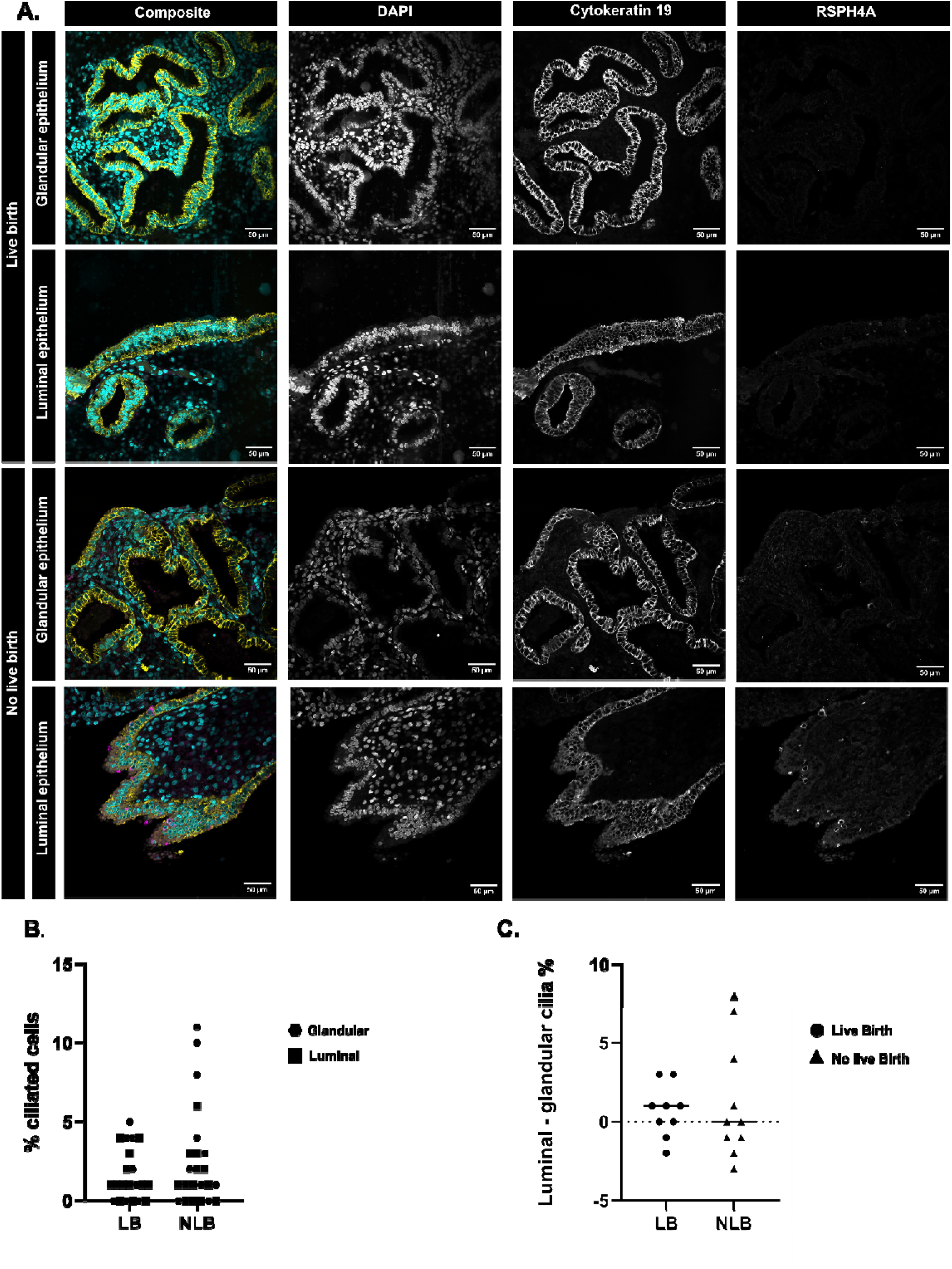
No differences in endometrial epithelial cell ciliation between those who had live birth and no live birth after endometrial scratch and IVF. **A**. Confocal micrographs of endometrial glandular and luminal epithelium at the window of implantation. DAPI staining nuclei (Cyan), Cytokeratin 19 staining epithelium (Yellow) and RSPH4A staining ciliated epithelial cells (Magenta). Scale bars = 50 µm. **B**. There is no significant difference between the percentage of ciliated cells between live birth and no live birth cohorts (p = 0.57, n = 44). **C**. No significant difference in the distribution of cilia on the luminal surface and the glandular surface between the two outcome cohorts (p = 0.82, n = 42)

## Discussion

In this study, we interrogated the endometrial phenotype at the window of implantation in a rigorously characterised IVF cohort following participation in a randomised trial of endometrial scratch. Using molecular dating to ensure cycle synchrony, bulk RNA sequencing to assess transcriptomic differences, and quantitative epithelial analysis to examine ciliation coverage, in endometrial biopsy samples, we found no significant differences between women who achieved live birth and those who did not. These findings suggest that, within a well-dated implantation window, neither global endometrial gene expression patterns nor endometrial epithelial ciliation coverage distinguishes subsequent pregnancy outcome in this population.

### Main findings of the study

First, we demonstrated concordance between urinary LH surge timing and transcriptomic dating algorithms, confirming that biopsies were accurately staged within the window of implantation. This is critical, as even small deviations in cycle timing can confound differential gene expression analyses. The absence of a temporal imbalance between live- and non-live-birth groups strengthens the validity of our negative findings.

Second, differential gene expression analysis revealed no significant transcriptomic differences between outcome groups when applying stringent thresholds for fold change and false discovery rate. Principal component analysis likewise showed no clustering by pregnancy outcome, indicating that the overall transcriptomic profile was comparable between groups. These data do not support the hypothesis that women who achieve live birth after IVF have a distinct global endometrial gene-expression signature at day LH+7, detectable by bulk RNA sequencing.

Third, automated quantification of endometrial epithelial cell ciliation coverage demonstrated no significant difference between groups. Although ciliated epithelial cells represent a hormonally responsive component of the endometrial luminal surface, our data suggest that absolute coverage at the window of implantation is not associated with clinical outcome in this setting.

### Strengths and Limitations of the study

A key strength of this study is the precise endometrial dating achieved through both urinary LH testing and transcriptomic modelling, reducing misclassification of menstrual cycle day within the implantation window. The cohort was derived from a well-defined, multicentre randomised trial population, limiting selection bias. Applying stringent thresholds for differential gene expression analysis and correcting for multiple comparisons reduces the likelihood of false-positive findings.

Automated epithelial cell quantification analysis minimised observer bias in structural assessment. Integration of molecular and histological analyses within the same biopsies provides a multidimensional evaluation of endometrial phenotype.

Several limitations merit consideration. Sample size, although adequate for detecting large transcriptional shifts, may be underpowered to detect small effect sizes. Bulk RNA sequencing cannot resolve cell-type-specific differences or spatial heterogeneity. Additionally, biopsy sampling represents a snapshot of a dynamic process; temporal fluctuations within the implantation window may not be captured at a single LH+7 time point.

Our cohort comprised women undergoing IVF within the context of a scratch trial, and findings may not extrapolate to women with recurrent implantation failure or other distinct populations. Finally, transcript abundance does not necessarily equate to protein expression or functional activity.

### Interpretation of findings in context of endometrial scratch literature

The biological rationale for endometrial scratching has centred on inducing a wound-healing inflammatory cascade, with downstream modulation of cytokines, adhesion molecules, and decidual pathways presumed to enhance receptivity. However, high-quality clinical trials in women undergoing first-time IVF have not demonstrated improvement in live birth rates. The absence of detectable endometrial transcriptomic differences in our cohort provides mechanistic support for these clinical findings.

Our data are consistent with recent longitudinal transcriptomic analyses showing no sustained alteration in peri-implantation endometrial gene expression following mid-luteal scratch in the preceding cycle. If scratch exerts any biological effect to the endometrium, it may be transient and confined to the immediate inflammatory phase rather than inducing durable remodelling detectable in the subsequent implantation window. Alternatively, its effects may be subtle, cell-type-specific, or post-transcriptional and therefore not captured by bulk RNA sequencing.

The discrepancy between neutral large, randomised trials and some meta-analyses suggesting modest benefit of ES may reflect population heterogeneity, variation in timing and technique of injury, or statistical aggregation of small effect sizes across diverse cohorts. Importantly, our study interrogates biological plausibility rather than clinical effect size and does not identify a molecular signature that would explain differential response to ES at the level of global gene expression.

### Endometrial transcriptomics and implantation biology

The concept of a discrete “receptive transcriptome” has emerged from bulk and single-cell profiling studies that describe gene-expression shifts during the window of implantation. However, receptivity is unlikely to be binary or solely transcriptionally defined. Implantation represents a complex, dynamic interplay between embryo competence and maternal environment. It is plausible that embryo-derived factors, stochastic processes, or microenvironmental gradients within the uterine cavity play a greater role than steady-state transcript abundance at a single time point.

Moreover, bulk RNA sequencing averages signals across heterogeneous cell populations. Subtle differences confined to rare epithelial or stromal subtypes, immune cell subsets, or spatial niches may not be detectable without single-cell or spatial transcriptomic approaches. Thus, the absence of differential expression in endometrium at the bulk level does not exclude more nuanced cellular variation.

### Epithelial ciliation and structural receptivity

Ciliated epithelial cells are traditionally associated with tubal gamete transport but are also present within the endometrium, where their functional significance remains incompletely defined. Hormonal regulation of ciliation suggests potential temporal roles in fluid dynamics and embryo apposition. However, our quantitative analysis did not identify differences in overall endometrial epithelium ciliation coverage between outcome groups. This suggests that the absolute proportion of endometrial ciliated epithelium at day LH+7 is unlikely to be a major determinant of implantation success in IVF.

It remains possible that functional aspects of epithelial cilia behaviour, such as beat frequency, localised distribution, or interaction with secretory cell populations, are more relevant than overall coverage in the endometrium. Future work integrating ultrastructural assessment and functional imaging of cilia may provide further insight into their role in the endometrium.

## Conclusion

Taken together, our findings suggest that, within a precisely timed implantation window, neither the global transcriptomic profile nor the epithelial ciliation coverage of the endometrium differentiates women who achieve live birth from those who do not. These data reinforce the concept that implantation failure after IVF is unlikely to be explained by major, detectable differences in steady-state endometrial gene expression or gross epithelial architecture alone.

In the context of endometrial scratch, our results do not provide mechanistic evidence supporting a sustained molecular effect in the subsequent cycle. Combined with neutral findings from large, randomised trials, this further calls into question the biological plausibility of routine scratching as a receptivity-enhancing intervention.

Future studies should employ single-cell and spatial transcriptomics to dissect cell-type-specific and microenvironmental variations. Integration of proteomics, immune profiling, and embryo-derived signalling analyses may better elucidate the maternal-embryo interface. Longitudinal sampling during the peri-implantation period may capture dynamic changes that are not apparent at a single time point.

Understanding implantation will likely require systems-level approaches rather than reliance on single-modality biomarkers.

## Authors Roles

Funding acquisition: J.C, Y.C, R.L, S.L, M.M; Conceptualisation: Y.C, J.C, M.M, E.P, S.L, R.L; Project administration: J.C, C.P; Supervision: J.C, Y.C, R.L, C.P; Methodology: S.L, C.P, J.FP, E.P, I.R; Investigation, Formal analysis, Data curation, Validation: J.FP, C.P, E.P, I.R; Writing – original draft preparation: E.P, Y.C; Writing – review & editing: All authors.

## Acknowledgements

We would like to thank the women who consented to be part of this study and clinical staff who assisted with sample collection. The authors acknowledge the use of the IRIDIS High Performance Computing Facility, and associated support services at the University of Southampton, in the completion of this work. The authors gratefully acknowledge the Faculty of Medicine BIO-R, at the University of Southampton for their support and assistance in this work.

## Funding

Funded by Wellbeing of Women (RG2147), Wessex Medical Research, and the Rosetrees Trust (PGS23/100171).

## Conflict of interest

No Conflicts of interest to declare

## Notes

### Competing Interest Statement

The authors have declared no competing interest.

